# Mutations in *ELAC2* associated with hypertrophic cardiomyopathy impair mitochondrial tRNA 3’-end processing

**DOI:** 10.1101/378448

**Authors:** Makenzie Saoura, Christopher A. Powell, Robert Kopajtich, Ahmad Alahmad, Haya H. AL-Balool, Buthaina Albash, Majid Alfadhel, Charlotte L. Alston, Enrico Bertini, Penelope Bonnen, Drago Bratkovic, Rosalba Carrozzo, Maria A. Donati, Michela Di Nottia, Daniele Ghezzi, Amy Goldstein, Eric Haan, Rita Horvath, Joanne Hughes, Federica Invernizzi, Eleonora Lamantea, Benjamin Lucas, Kyla-Gaye Pinnock, Maria Pujantell, Shamima Rahman, Pedro Rebelo-Guiomar, Saikat Santra, Daniela Verrigni, Robert McFarland, Holger Prokisch, Robert W. Taylor, Louis Levinger, Michal Minczuk

**Affiliations:** York College/CUNY, Jamaica, NY 11451, USA; MRC Mitochondrial Biology Unit, University of Cambridge, Cambridge CB2 0XY, UK; Institute of Human Genetics, Technische Universität München, 81675 Munich, Germany; Institute of Human Genetics, Helmholtz Zentrum München, 85764 Neuherberg, Germany; Wellcome Centre for Mitochondrial Research, Institute of Neuroscience, Newcastle University, Newcastle upon Tyne NE2 4HH, UK; Kuwait Medical Genetics Center, Ghanima Alghanim building, Al-Sabah Medical Area, Kuwait; Genetics Division, Department of Pediatrics, King Saud bin Abdulaziz University for Health Sciences, King Abdullah International Medical Research Centre, King Abdulaziz Medical City, MNG-HA, Riyadh, Saudi Arabia; Department of Neurosciences, Unit of Muscular and Neurodegenerative Disorders, Laboratory of Molecular Medicine, Bambino Gesu’ Children’s Research Hospital, 00165 Rome, Italy; Department of Molecular and Human Genetics, Baylor College of Medicine, Houston, TX 77030, USA; Adult Genetics Unit, Royal Adelaide Hospital, Adelaide SA 5000 and School of Health and Medical Sciences, The University of Adelaide, Adelaide SA 5005, Australia; Metabolic Unit, A. Meyer Children’s Hospital, 50139 Florence, Italy; Unit of Medical Genetics and Neurogenetics, Foundation IRCCS Institute of Neurology “Besta”, 20126 Milan, Italy; Department of Pathophysiology and Transplantation, University of Milan, Milan, Italy; Mitochondrial Medicine Frontier Program, Children’s Hospital of Philadelphia; Wellcome Centre for Mitochondrial Research, Institute of Genetic Medicine, Newcastle University, Newcastle upon Tyne NE1 3BZ, UK; National Centre for Inherited Metabolic Disorders, Temple Street Children’s University Hospital, Dublin, Ireland; Mitochondrial Research Group, UCL Great Ormond Street Institute of Child Health, London WC1N 1EH, UK; Graduate Program in Areas of Basic and Applied Biology, University of Porto, Porto, 4200-135, Portugal; Department of Clinical Inherited Metabolic Disorders, Birmingham Children’s Hospital, Steelhouse Lane, Birmingham, UK

## Abstract

Dysfunction of mitochondrial gene expression, caused by mutations in either the mitochondrial or nuclear genomes, is associated with a diverse group of human disorders characterized by impaired mitochondrial respiration. Within this group, an increasing number of mutations have been identified in nuclear genes involved in mitochondrial RNA metabolism. For instance, pathogenic mutations have been identified in the genes encoding enzymes involved in the precursor transcript processing, including *ELAC2.* The *ELAC2* gene codes for the mitochondrial RNase Z, which is responsible for endonucleolytic cleavage of the 3’ ends of mitochondrial pre-tRNAs. Here, we report the identification of sixteen novel *ELAC2* variants in individuals presenting with mitochondrial respiratory chain deficiency, hypertrophic cardiomyopathy and lactic acidosis. We provided further evidence for the pathogenicity of the three previously reported variants by studying the RNase Z activity in an *in vitro* system and applied this recombinant system to investigate all novel missense variants, confirming the pathogenic role of these new *ELAC2* mutations. We also modelled the residues affected by missense mutation in solved RNase Z structures, providing insight into enzyme structure and function. Finally, we show that primary fibroblasts from the individuals with novel *ELAC2* variants have elevated levels of unprocessed mitochondrial RNA precursors. Our study thus broadly confirms the correlation of *ELAC2* variants with severe infantile-onset forms of hypertrophic cardiomyopathy and mitochondrial respiratory chain dysfunction. One rare missense variant associated with the occurrence of prostate cancer (p.Arg781His) impairs the mitochondrial RNase Z activity of ELAC2, possibly indicating a functional link between tumorigenesis and mitochondrial RNA metabolism.

## Introduction

Mitochondria are essential for cell function through their involvement in ATP synthesis by oxidative phosphorylation (OXPHOS), and host a number of other biosynthetic pathways. Whereas the majority of the mitochondrial proteome is encoded in the cell nucleus and imported into the mitochondria following synthesis on cytosolic ribosomes, 13 essential subunits of the OXPHOS system are synthesised within the organelle. For this reason, mitochondria maintain and express mitochondrial DNA (mtDNA) that encodes the aforementioned polypeptides, together with the mitochondrial (mt-) tRNAs and rRNAs. All remaining protein components of the mitochondrial gene maintenance and expression machinery such as proteins responsible for mtDNA transcription, precursor RNA processing enzymes, the mitoribosomal proteins, mitochondrial aminoacyl tRNA synthetases and others are encoded by the nuclear genes [1, 2]. More than 50 nuclear-encoded mitochondrial proteins involved in mitochondrial gene expression are linked to heritable disorders [3–7].

Human mt-mRNAs, mt-tRNAs, and mt-rRNAs are transcribed as part of large polycistronic precursor transcripts encoded by the 16.6 kb mtDNA. The mitochondrial gene sequences for mt-rRNA and, in the majority of cases, mt-mRNA are separated by mt-tRNA sequences, leading to the proposed ‘tRNA punctuation’ model of RNA processing [8, 9]. Processing of the intervening mt-tRNAs generates individual mt-mRNAs and mt-rRNAs. In human mitochondria, endonucleolytic cleavage at the 5′ and 3′ termini of mt-tRNAs is performed by the RNase P complex [10] and the RNase Z activity of ELAC2, respectively [11]. Two human nuclear genes encode orthologues of the bacterial RNase Z (elaC): *ELAC1* (MIM 608079, RefSeq: NM_0018696.2, NP_061166.1) and *ELAC2* (MIM 605367, RefSeq: NM_018127.6, NP_060597.4). The *ELAC1* gene encodes a short form of RNase Z (also referred to as RNase Z^S^) that is located in the cytosol, but its function in humans is unknown. *ELAC2* codes for a long form of RNase Z (also referred to as RNase Z^L^). Alternative translation initiation of *ELAC2* mRNA has been proposed to produce two ELAC2 protein isoforms; one targeted to the mitochondria, the other to the nucleus. Translation of the longer isoform initiates at the methionine codon 1 (M1) and the first 31 amino acids of this isoform are predicted to contain a mitochondrial targeting sequence (MTS). This longer isoform of ELAC2 has a well-characterized role in mitochondrial pre-tRNA processing [11–14]. If translation is initiated at the methionine codon 16 (M16) (relative to the longer mitochondrially-targeted form) ELAC2 is targeted to the nucleus, where it is predicted to be responsible for nuclear pre-tRNA 3’ end maturation [11].

Thus far, six different pathogenic mutations have been reported in *ELAC2*-associated mitochondrial dysfunction. In our previous work, we investigated the functional consequences of pathogenic *ELAC2* variants coding for p.Phe154Leu, p.Arg211, p.Leu423Phe and p.Thr520Ile in patients presenting with a recessively inherited form of hypertrophic cardiomyopathy (HCM), hypotonia, lactic acidosis, and failure to thrive [15]. This previous work identified a total of four disease alleles (three missense and one nonsense) in three families. One of these, the variant coding for p.Phe154Leu, was also recently reported as prevalent in consanguineous Arabian families affected by infantile cardiomyopathy [16]. In contrast, a homozygous splice site mutation (c.1423+2T>A) in *ELAC2* has been associated with developmental delay and minimal cardiac involvement in a consanguineous Pakistani family [17]. Finally, a single heterozygous *ELAC2* variant coding for p.Pro32Arg was recently reported in an infant presenting with encephalopathy, epilepsy, and growth and developmental retardation [18]. The patient also had a cardiac anomaly, however, without evidence of cardiomyopathy. Since the transmission pattern of *ELAC2*-related disease in the families reported previously [15–17] was consistent with recessive inheritance, the relevance of this variant remains unclear.

In the present work, we report the identification of sixteen additional *ELAC2* variants (ten missense, two frameshift and four splice mutations) in individuals who present with mitochondrial respiratory chain deficiency, HCM and lactic acidosis. We provide further evidence for the pathogenicity of the three previously reported and ten newly identified missense variants by studying the RNase Z activity in an *in vitro* system. Moreover, fibroblasts from the individuals with novel *ELAC2* variants showed elevated levels of unprocessed mt-tRNA precursors. Modelling of the missense substitutions provided additional insight into the effects of substitutions on enzyme structure.

## Methods

### Ethics statement

Informed consent for diagnostic and research-based studies was obtained for all subjects in accordance with the Declaration of Helsinki protocols and approved by local institutional review boards.

### Exome Sequencing, variant prioritization, reevaluation and verification

For reevaluation of the *ELAC2* variants in P1 (previously reported as patient 27 in [19]) and also for the analysis of the *ELAC2* splice variants in P6 and P8, RNA purification from fibroblasts and cDNA retrotranscription was used to verify the identified variants, we used RNeasy mini kit (QIAGEN) and GoTaq 2-Step RTqPCR System (Promega), respectively, according to the manufacturers’ protocols. Primers used for cDNA amplification are available upon request.

For P2 and P10-P13 exome sequencing and variant prioritisation was performed by commercial laboratories: P2 - Fulgent Diagnostics, P10 and P12 - Baylor College of Medicine, Human Genome Sequencing Center, P11-Centogene, as described in previously [20] and P13 – UCL.

For P3, targeted NGS sequencing using a custom Ampliseq panel targeting 55 mitochondrial translation genes (IAD62266) and subsequent Ion Torrent PGM sequencing was performed essentially as described previously [21]. Candidate gene sequencing was performed for all coding exons of the *ELAC2* gene (including intron-exon boundaries) using M13-tagged amplicons and BigDye v3.1 sequencing kit (Life Technologies). Capillary electrophoresis was performed using an ABI3130xl (Life Technologies). NGS variant confirmation was performed by Sanger sequencing using oligonucleotides targeting the exons of interest.

For P4 and P8, genomic DNA from the individuals and their parents was isolated from whole blood using the chemagic DNA Blood Kit special (PerkinElmer,Waltham, USA), according to the manufacturer’s protocol. Exome sequencing was performed as previously described [22]. Exonic regions were enriched using the SureSelect Human All Exon kit (50Mb_v5) from Agilent followed by sequencing as 100 bp paired-end runs on an Illumina HiSeq2500. Reads were aligned to the human reference genome (UCSC Genome Browser build hg19) using Burrows-Wheeler Aligner (v.0.7.5a). Identification of single-nucleotide variants and small insertions and deletions (indels) was performed with SAMtools (version 0.1.19). For analysis of rare bi-allelic variants, only variants with a minor allele frequency (MAF) of less than 1% in our internal Munich database of 14,000 exomes were considered.

For P5, WES was undertaken using previously described methodologies and bioinformatics variant filtering pipelines [23]

For P6 and P9 a targeted custom panel (Nextera rapid capture, Illumina) containing genes responsible for mitochondrial disorders was used [24]. Variant filtering was performed as described in Legati *et al.* [25].

For P7, exome capture and massively parallel sequencing were outsourced (BGI, Shenzhen, China) using Sure Select Human All Exon V.4 Agilent and deep Illumina HiSeq technology (median reads depth = 50×). High quality variants were filtered against public (dbSNP146 and EXAC V.0.3) and in-house database, to retain private, rare (MAF less than 1%) and clinically pathogenic nucleotide changes. Variants prioritisation in the context of their functional impact was performed using *in silico* programs for missense mutations (Sift and Poliphen2) and also taking into account changes potentially affecting splice sites. All variants identified by NGS were validated by Sanger Sequencing as well the segregation in the family.

### RNA isolation and RNA northern blotting

RNA extraction and northern blotting were performed essentially as described previously [26]. Briefly, RNA was extracted from cells at 60 - 80 % confluency using TRIzol reagent (Ambion), following the manufacturer’s instructions. Gels run on the Bio-Rad Mini-Sub Cell GT were used for the separation of RNA samples (5 µg per sample) for northern blotting. A half volume of Gel Loading Buffer II (Ambion) was added to samples before heating at 55 °C for 10 minutes, chilled on ice for 2 minutes, and then loaded onto a 1.2 % Agarose gel (1 x MOPS, 0.7 M formaldehyde). Electrophoresis was carried out at 4 °C in 1 x MOPS, 0.3 M formaldehyde and 10 µg/mL ethidium bromide. Gels were semi-dry blotted onto a positively charged nylon membrane (Hybond-N+, GE Healthcare) for >12 hours, after which the membrane was cross-linked and hybridized with [^32^P] labelled antisense RNA probes as described previously [27].

### Recombinant protein purification and mutagenesis

Wild type ELAC2 protein (GenBank Accession number: NM_018127.6) was expressed from Gly50 to Gln826 using the baculovirus system and insect SF9 cells, and affinity purified as previously described [28]. Missense mutants were constructed by overlap extension PCR. Subcloning sites used for substitutions in the amino domain and linker were introduced the *Bam*HI site at the amino end to the internal natural *Eco*RI site (nt 1450 in NM_018127) and in the carboxy domain from the *Eco*RI site to the introduced *Xho*I site following the termination codon. Sequences of the mutant constructs were confirmed by Sanger sequencing (Macrogen).

### RNase Z processing experiments

The mitochondrial pre-tRNA substrates for RNase Z reactions were prepared by runoff T7 transcription using cis-acting hammerheads to cleave at +1. The mt-tRNA^Leu(UUR)^ substrate has a 29 nt 3’-end trailer with natural sequence ending with a *Sma*I runoff (-CCC) and mt-tRNA^Ile^ has a 19 nt 3’-trailer, also ending with a *Sma*I runoff. Substrate 5’ ends were radiolabelled using [γ-^32^P]-ATP and polynucleotide kinase. For processing experiments, the concentration of unlabelled substrate was varied, over the range from 4 – 100 nM, at a constant much lower concentration of labelled substrate used as a tracer. The enzyme was used at the lowest concentration that produces a quantifiable product band at the highest concentration of unlabelled substrate used in the experiment. Kinetic experiments were performed with wild type enzyme at 10 pM using mt-tRNA^Leu(UUR)^ substrate, and at 50 pM using mt-tRNA^Ile^, as in previous experiments [29]. Mutant enzymes were used at a higher concentration than wild type depending on the impairment factor (**Figure S1**). Variant processing experiments were performed in parallel with wild-type on the same day. Reactions were performed using Processing Buffer consisting of 25 mM Tris-Cl pH 7.2, 1.5 mM CaCl_2_, 1 mM freshly prepared Dithiothreitol and 0.1 mg/ml BSA. Reactions were sampled after 5, 10 and 15 min incubation at 37°C, electrophoresed on denaturing 6% polyacrylamide gels and images were obtained from dried gels using a storage phosphor screen and Typhoon scanner and analysed with ImageQuant TL (GE Life Sciences). In this design with constant trace labelled substrate and varying concentration of unlabelled substrate the measurement of proportion of product per minute of reaction is equivalent to V/[S]. V is obtained by multiplying by [unlabelled S] for each reaction, as illustrated in the processing data figures.

### In silico modelling

The recently published structure of *Saccharomyces cerevisia*e RNase Z (PDB 5MTZ [30]) was used to model the position and function of residues at which substitutions were observed in this study. Molecular structures were displayed using PyMOL. Overall structure is shown using cartoon and individual residues with sticks, polar contacts with dashed lines and hydrophobic residues with dots. The active site and regions proximal to it, which directly contact the substrate, are found in the carboxy domain of ELAC2 (a long form RNase Z, RNase Z^L^), were superimposed onto the corresponding regions in the only available co-crystal structure of RNase Z with pre-tRNA substrate, from *Bacillus subtilis* (PDB 4GCW) [31].

## Results

### Summary of clinical features of the investigated patient cohort

We investigated 13 families with a cohort of 13 infants, most presenting with early-onset, progressive cardiomyopathy, with hypertrophic cardiomyopathy (HCM) being present in 10 subjects, dilated cardiomyopathy (DCM) in 2 subjects and one subject being reported without any cardiac problems. All subjects with the exception of P10 also presented with lactic acidosis. These clinical features raised suspicion of mitochondrial disease. Indeed, a biochemical defect of the mitochondrial respiratory chain (MRC) complexes was detected in all investigated subjects (n=10), with isolated Complex I deficiency prevailing in most of the patients (7/10) and the remaining subjects presenting with combined MRC deficiencies. The onset of symptoms was either from birth (P1, P2 and P11), neonatal (P5), infantile (P4, P6-P10, P12 and P13) or early childhood (P3). Most of the patients also displayed developmental delay. Evidence of brain involvement was found in P2, P5 and P9. P7 and P10 were successfully treated by heart transplantation at age 3.8 years and 10 months, respectively [32, 33], whereas P12 underwent two failed cardiac transplants. The summary of genetic, biochemical, and clinical findings of all individuals is provided in **Table 1**. Pedigrees of investigated families and detailed case reports are provided in the **Supplementary Information**.

**Table 1.** Patient summary

| Case No | cDNA (NM_018127.6) | Protein (NP_060597) | Identification | Sex | Age-at-onset | Cardiomyopathy | MRC deficiency | Additional clinical features | Previously reported |
| --- | --- | --- | --- | --- | --- | --- | --- | --- | --- |
| #57415 | c.631C>T; c.1559C>T | p.Arg211*; p.Thr520Ile | WES | male | 3 months | HCM | CI | Psychomotor retardation, mild hypotonia, lactic acidosis, sensorineural hearing impairment, hyperintensities in basal ganglia at age 3 m | [15] |
| #61982 | c.460T>C; c.460T>C | p.Phe154Leu; p.Phe154Leu | WES | female | 2 months | HCM | CI | Intrauterine growth retardation, lactic acidosis, cardiac failure; normal muscle biopsy findings | [15] |
| #65937 | c.1267C>T; c.1267C>T | p.Leu423Phe; p.Leu423Phe | WES | female | 5 months | HCM | CI | Psychomotor retardation, muscular hypotonia, cardiac failure | [15] |
| P1 | c.1478C>T; c.202C>T | p.Pro493Leu; p.Arg68Trp | WES | female | Birth | HCM | CI+CIV | Lactic acidosis, muscle weakness, ragged-red fibres and COX-deficient/SDH- hyper-reactive fibres. | [34] |
| P2 | c.2009del; c.1423+1G>A | p.Cys670Serfs*14; CSM | GP | male | Birth | No | ND | Lactic acidosis, developmental delay, ataxia, microcephaly, constipation, cerebellar vermis hypoplasia and prominent posterior fossa on brain.MRI | this report |
| P3 | c.297-2_297delinsTG; c.2342G>A | CSM; p.Arg781His | GP | female | 18 months | HCM | CI | Elevated blood lactate level (normal serum levels), developmental delay, IUGR | this report |
| P4 | c.2186A>G; c.2342G>A | p.Tyr729Cys; p.Arg781His | WES | female | 2 months | HCM | CI | Lactic acidosis, cardiovascular collapse | this report |
| P5 | c.460T>C; c.460T>C | p.Phe154Leu; p.Phe154Leu | WES | male | neonatal | HCM | ND | Lactic acidosis, fatal infantile cardiomyopathy | this report |
| P6 | c.798-1G>T; c.1690C>A | CSM; p.Arg564Ser; | Gene panel | female | 4 months | DCM | CI | Lactic acidosis, developmental delay, fatal infantile cardiomyopathy, EF 30% | this report |
| P7 | c.1979A>T; c.2039C>T | p.Lys660Ile; p.Ala680Val; | WES | female | 12 months | HCM | CI+CIV | Elevated blood lactate level, Patient transplanted at age 3.8 years | [31] |
| P8 | c.245+2T>A; c.1264C>G | CSM; p.Leu422Val | WES | female | 2 months | DCM | CI | Lactic acidosis, fatal infantile cardiomyopathy, EF 20-30% | this report |
| P9 | c.1163A>G; c.1163A>G | p.Gln388Arg; p.Gln388Arg | GP | male | 6 months | HCM | CI | Lactic acidosis, psychomotor retardation, fatigability, peripheral neuropathy. | this report |
| P10 | c.457delA; c.2342G>A | p.Ile153Tyrfs*6; p.Arg781His | WES | male | 8 months | HCM | CI | Developmental delay, hypotonia, GI dysmotility, s/p cardiac transplant | [32] |
| P11 | c.460T>C; c.460T>C | p.Phe154Leu; p.Phe154Leu | WES | female | Birth | HCM | CI | Lactic acidosis, later mild muscular hypotonia, lipid storage myopathy on skeletal muscle biopsy | this report |
| P12 | c.297-2_297-1delinsT; c.2245C>T | CSM; p.His749Tyr | WES | female | 4 months | HCM | CI+CIV | Lactic acidosis, global developmental delay, hypotonia, s/p cardiac transplant | this report |
| P13 | c.460T>C; c.460T>C | p.Phe154Leu; p.Phe154Leu | WES | female | 5 months | HCM | ND | Lactic acidosis, failure to thrive, | this report |

### Identification of ELAC2 variants

Using whole exome sequencing (WES) or targeted, panel-based next-generation sequencing, we screened a group of patients presenting with MRC deficiencies and cardiomyopathy (**Table 1**). Three of the analyzed patients (P5, P11 and P13) harbored the previously described homozygous c.460T>C (p.Phe154Leu) *ELAC2* variant [15, 16]. All remaining patients of the analyzed cohort harbored novel missense (n=10), frameshift (n=2) or splice site (n=4) mutations (**Figure 1** and **Table 1**). P2 harbored a novel consensus splice variant (c.1423+1G>A) in the same splice site as previously reported patients of Pakistani origin (c.1423+2T>A) [17]. All these variants were extremely rare or not reported in public databases and predicted damaging using Polyphen-2 [34].

**Figure 1.**
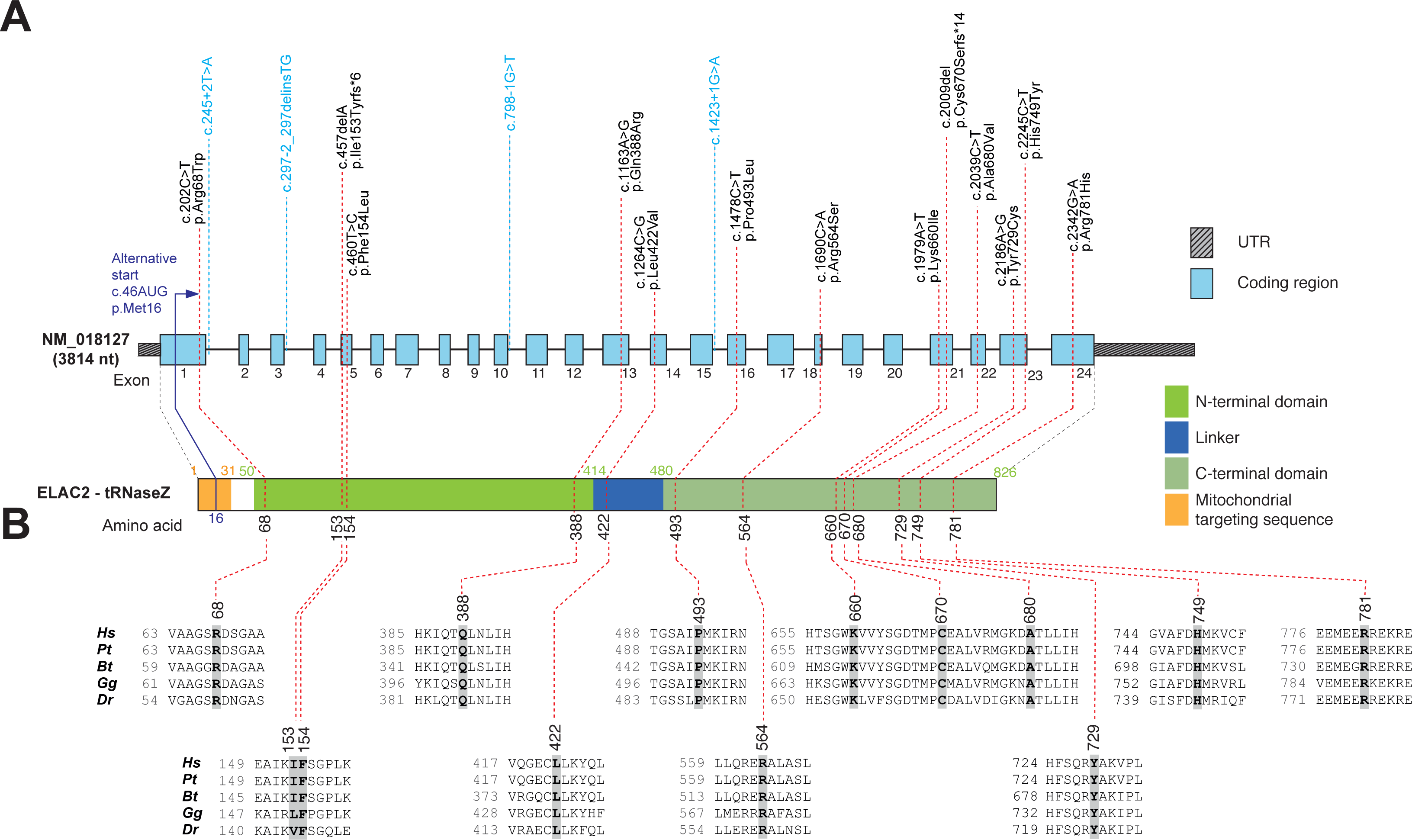
ELAC2 mutations and gene/protein structure. (A) Gene structure of *ELAC2* with known protein domains (as defined in [35]) of the gene product and localization of amino acid residues and splice sites (variants indicated in blue) affected by mutations. Intronic regions are not drawn to scale. **(B)** Conservation of human ELAC2 amino acid residues affected by mutations across *Pongo tapanuliensis* (Pt), *Bos Taurus* (Bt), *Gallus gallus* (Gg) and *Danio rerio* (Dr). For complete sequence alignment including *Saccharomyces cerevisiae* Trz1 used for structure modelling, see **Figure S3**.

Of note, P1 was previously included in WES analysis based on multiple MRC complex deficiency together with a cohort of 53 patients (patient 27) [19]. This previous study identified heterozygous c.1478C>T (p.Pro493Leu) and c.1621G>A (p.Ala541Thr) *ELAC2* variants and predicted them to be pathogenic. Re-evaluation of the c.1621G>A (p.Ala541Thr) variant revealed a minor allele frequency of 3.4% in the gnomAD database, with 218 homozygote individuals, which precludes pathogenicity and prompted analysis of cDNA obtained from P1 fibroblasts; this revealed that the c.1478C>T (p.Pro493Leu) and c.1621G>A (p.Ala541Thr) variants are situated *in cis*. Further, analysis of the cDNA sequence revealed a heterozygous change c.202C>T (p.Arg68Trp), which was absent in the gnomAD database and affects a moderately conserved amino acid. The c.202C>T substitution, located in a region which was not covered by the previous WES, was confirmed also in genomic DNA of P1.

*ELAC2* transcript analysis performed in the other two patients with splice site variants confirmed their deleterious effects. In P6 (compound heterozygous c.1690C>A; c.798-1G>T), we found mono-allelic expression of c.1690A, and no aberrant species, indicating that the splice variant causes mRNA decay. In P8 (compound heterozygous c.245+2T>A; c.1264C>G) we amplified two species of *ELAC2* transcripts: the full-length and one missing exon 1.

### In vitro RNase Z activity of mutant ELAC2 enzymes

Despite the increased utility of genetic testing, providing proof of pathogenicity of novel variants remains challenging and follow up functional studies *in vitro* should therefore be included as an integral part of the evaluation. In order to provide evidence for the pathogenicity of identified *ELAC2* variants, we set out to study the RNase Z activity of the enzyme in the presence of the missense substitutions or the truncating variants resulting from frameshift mutations. Substitutions were introduced into the human *ELAC2* cDNA (Genbank Acc# NM_018127.6) and the mutant proteins were expressed in baculovirus using insect SF9 cells [35]. Affinity-purified recombinant mutant proteins were tested using precursor mt-tRNA^LeuUUR^ or mt-tRNA^Ile^ as substrates.

In order to assess the utility of the *in vitro* system to evaluate the pathogenicity of the novel *ELAC2* variants, we first tested the three previously reported missense mutations (p.Phe154Leu, p.Leu423Phe, p.Thr520Ile), that have been extensively characterized in terms of pathogenicity in our previous paper [15]. Recombinant mutant proteins expressed well (**Figure S1a**), suggesting that the substitutions do not severely alter stability of ELAC2. Next, we obtained Michaelis-Menten plots and kinetic parameters in comparative kinetic experiments testing endonucleolytic cleavage of mitochondrial pre-tRNA using wild-type and mutant ELAC2 preparations (**Table 2, Figure 2a** and **Figure S1b-c**). All three mutants tested showed significant impairment of the RNAse Z activity, exhibiting reduced *k*_cat_/*K*_M_ values as compared to the wild-type enzyme (**Figure 2a**). The impairment observed for the mutant proteins was largely due to reduced *k*_cat_. In these three instances the impairment is moderate, with *k*_cat_/*K*_M_ being in the range of 20–80% of the WT enzyme, consistent with the essential role of ELAC2 in mt-tRNA processing. This result further confirms the pathogenicity of the p.Phe154Leu, p.Leu423Phe, p.Thr520Ile variants and is consistent with the previously reported accumulation of mtRNA precursors of 3′-end unprocessed mt-tRNA in patient-derived cells [15]. Importantly, these data also establish the utility of the assay for testing newly-detected *ELAC2* variants.

**Table 2.** Kinetic Parameters of three previously reported pathogenic missense mutations in the ELAC2 endonuclease with the pre-mt-tRNA^Leu(UUR)^ substrate. Column designation, from left: **(column 1)** Variant: wild-type (WT) or mutant ELAC2. **(column 2)** number of times variant processing experiments were performed. **(columns 3-5)** *k_cat_*, *KM*, *k_cat_*/*K_M_*: values reported are means of replicate experiments. Values following ± are standard errors. **(columns 6-8)** *k_cat_*, *K_M_* and *kcat*/*KM* relative to WT (e.g. the quotient [*k*_cat_mutant]/[*k*_cat_ WT]). *Reported variant relative to WT values are the means and standard errors for specific experiments performed on the same day, rather than the compiled values at the top of the table, therefore differ from results that would be obtained by comparing values in columns to the left with aggregate means for WT (first row).

| Mutant | # Trials | $k_{cat}$<br>(min<sup>-1</sup>) | $K_M$<br>(nM) | $k_{cat}/K_M$<br>(x 10<sup>8</sup> M<sup>-1</sup>min<sup>-1</sup>) | Relative<br>$k_{cat}$ * | Relative<br>$K_M$ * | Relative<br>$k_{cat}/K_M$ * |
| --- | --- | --- | --- | --- | --- | --- | --- |
| WT | 11 | 23.7±2.8 | 78±8.0 | 3.2±0.36 |  |  |  |
| Phe154Leu | 4 | 12±2.80 | 140±36 | 0.84±0.12 | 0.51±0.06 | 1.9±0.53 | 0.32±0.06 |
| Leu423Phe | 3 | 28±12 | 79±24 | 3.3±0.45 | 0.94±0.11 | 1.2±0.19 | 0.79±0.06 |
| Thr520Ile | 4 | 7.8±0.51 | 8.1±0.19 | 0.97±0.04 | 0.39±0.06 | 1.1±0.16 | 0.39±0.05 |

**Figure 2.**
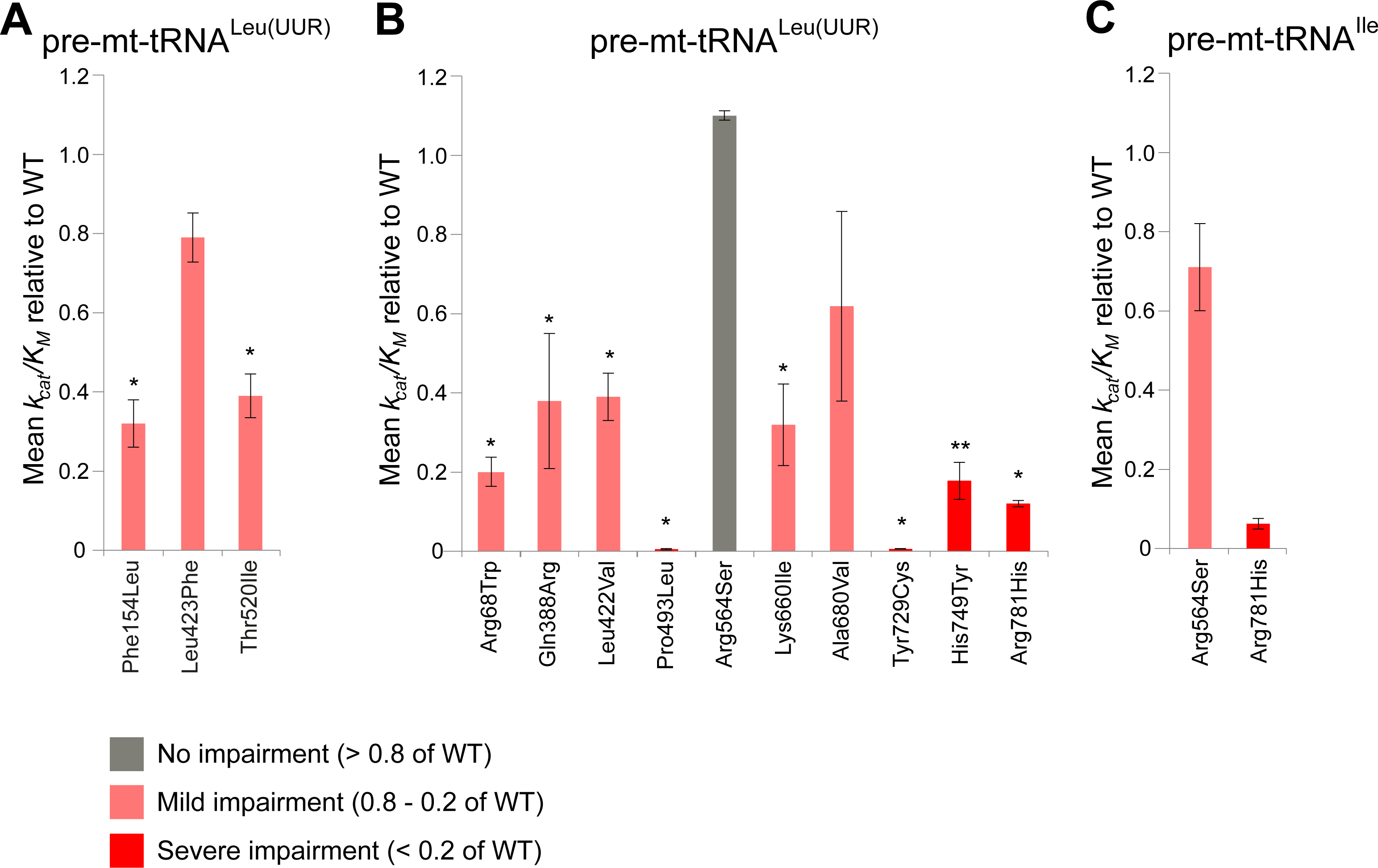
Kinetic parameters of the pathogenic ELAC2 variants. **(A)** Kinetic parameters of the previously reported ELAC2 variants [15] with mt-tRNALeu(UUR) substrate. The graph shows *k*_cat_/*K*_M_value relative to WT (e.g. the quotient [*k*_cat_ mutant]/[*k*_cat_WT]). The bars indicate standard error. *, ** above the bars indicate p-values with a significance of 0.05 and 0.01, respectively, calculated using unpaired t-test. **(B)** Kinetic parameters of the novel ELAC2 variants with mt-tRNA^Leu(UUR)^ substrate analysed as per (A). **(C)** Kinetic parameters of selected novel ELAC2 variants with mt-tRNA^Ile^ substrate analysed as per (A).

Next, we expressed and tested 10 novel missense variants detected in our patients using pre-mt-tRNA^Leu(UUR)^ as a substrate (**Table 3, Figure 2b**). A severe reduction of *k*_cat_/*K*_M_ values (below 20% of the wild type, WT, enzyme) was observed for p.Pro493Leu, p.Tyr729Cys, p.His749Tyr and p.Arg781His. Moderate impairment of the *k*_cat_/*K*_M_ values (20-80% of the WT enzyme) was observed for p.Arg68Trp, p.Gln388Arg, p.Leu422Val and p.Lys660Ile. The *k*_cat_/*K*_M_ values for p.Arg564Ser was not significantly affected for the pre-mt-tRNA^Leu(UUR)^ substrate. Reduced *k*_cat_ is principally responsible for impairment of mutant enzymes, however, a substantial increase in *K*_M_ relative to WT was observed for p.Arg781His; additionally the p.Pro493Leu and p.Tyr729Cys mutants exhibited lesser increases in *K*_M_ (**Table 3**). Previous data from RNASeq and northern blots indicated junction-dependent impairment of endonucleolytic cleavage of pre-tRNA by mutant ELAC2 enzymes [15]. With this in mind, we tested the p.Arg564Ser mutant (which did not show detectable impairment with the pre-mt-tRNA^Leu(UUR)^) using a different substate, pre-mt-tRNA^Ile^. Moderate impairment of the *k*_cat_/*K*_M_ values ratios (in the range of 20-80% of the WT enzyme) was observed for p.Arg564Ser with the pre-mt-tRNA^Ile^ substrate (**Figure 2c**, **Table 4**).

**Table 3.** Kinetic Parameters of the novel missense mutations in the ELAC2 endonuclease with the pre-mt-tRNA^Leu(UUR)^ substrate. Column designation, from left: **(column 1)** Variant: wild-type (WT) or mutant ELAC2. **(column 2)** number of times variant processing experiments were performed. **(columns 3-5)** *k_cat_*, *K_M_*, *k_cat_*/*K_M_*: values reported are means of replicate experiments. Values following ± are standard errors. **(columns 6-8)** *k_cat_*, *K_M_* and *k_cat_*/*K_M_* relative to WT (e.g. the quotient [*k*_cat_ mutant]/[*k*_cat_ WT]). *Reported variant relative to WT values are the means and standard errors for specific experiments performed on the same day, rather than the compiled values at the top of the table, therefore differ from results that would be obtained by comparing values in columns to the left with aggregate means for WT (first row). Grey – no impairment of enzymatic activity (> 0.8 of WT), light red – mild impairment of enzymatic activity (0.8 - 0.2 of WT), red – severe impairment of enzymatic activity (< 0.2 of WT).

| Mutant | # Trials | $k_{cat}$<br>(min<sup>-1</sup>) | $K_M$<br>(nM) | $k_{cat}/K_M$<br>(x 10<sup>8</sup> M<sup>-1</sup>min<sup>-1</sup>) | Relative<br>$k_{cat}$ | Relative<br>$K_M$ | Relative<br>$k_{cat}/K_M^*$ |
| --- | --- | --- | --- | --- | --- | --- | --- |
| WT | 34 | 20.2±2.1 | 43±7.7 | 6.2±0.56 |  |  |  |
| Arg68Trp | 4 | 1.9±0.36 | 37±16 | 0.81±0.22 | 0.13±0.032 | 0.79±0.25 | 0.20±0.038 |
| Gln388Arg | 4 | 1.2±0.46 | 14±7.8 | 1.5±0.66 | 0.075±0.023 | 0.43±0.18 | 0.38±0.17 |
| Leu422Val | 3 | 9.1 ±3.5 | 30±15 | 4.0±1.5 | 0.59±0.081 | 1.6±0.39 | 0.39±0.060 |
| Pro493Leu | 4 | 0.09±0.02 | 99±51 | 0.014±0.0035 | 0.006±0.002 | 2.0±0.8 | 0.005±0.002 |
| Arg564Ser | 4 | 44.1±19.3 | 40±27 | 11±2.8 | 2.2±0.68 | 2.5±1.3 | 1.1±0.16 |
| Lys660Ile | 5 | 3.2±1.4 | 25±10 | 1.5±0.23 | 0.23±0.12 | 1.2±0.54 | 0.32±0.10 |
| Ala680Val | 4 | 4.8±1.6 | 14±4.8 | 4.0±1.2 | 0.23±0.05 | 0.59±0.22 | 0.62±0.24 |
| Tyr729Cys | 3 | 0.1±0.01 | 23±8.2 | 0.056±0.022 | 0.006±0.001 | 0.77±0.26 | 0.0075±0.0015 |
| His749Tyr | 6 | 1.1±0.12 | 23±8.2 | 1.1±0.23 | 0.06±0.01 | 0.45±0.15 | 0.18±0.044 |
| Arg781His | 3 | 7.6±0.70 | 131±24 | 0.63±0.14 | 0.37±0.17 | 2.9±1.1 | 0.12±0.013 |

**Table 4.** Kinetic Parameters of the selected novel missense mutations in the ELAC2 endonuclease with the pre-mt-tRNA^Ile^ substrate. Column designation, from left: **(column 1)** Variant: wild-type (WT) or mutant ELAC2. **(column 2)** number of times variant processing experiments were performed. **(columns 3-5)** *k_cat_*, *K_M_*, *k_cat_*/*K_M_*: values reported are means of replicate experiments. Values following ± are standard errors. **(columns 6-8)** *k_cat_*, *K_M_* and *k_cat_*/*K_M_* relative to WT (e.g. the quotient [*k*_cat_ mutant]/[*k*_cat_ WT]). *Reported variant relative to WT values are the means and standard errors for specific experiments performed on the same day, rather than the compiled values at the top of the table, therefore differ from results that would be obtained by comparing values in columns to the left with aggregate means for WT (first row). Light red – mild impairment of enzymatic activity (0.8 - 0.2 of WT), red – severe impairment of enzymatic activity (< 0.2 of WT).

| **Mutant** | **# Trials** | **$k_{cat}$**<br>**(min<sup>-1</sup>)** | **$K_M$**<br>**(nM)** | **$k_{cat}/K_M$**<br>**(x 10<sup>8</sup> M<sup>-1</sup>min<sup>-1</sup>)** | **Relative**<br>**$k_{cat}$** | **Relative**<br>**$K_M$** | **Relative**<br>**$k_{cat}/K_M^*$** |
| --- | --- | --- | --- | --- | --- | --- | --- |
| **WT** | 3 | 0.17±0.05 | 3.4±1.3 | 0.57±0.17 |  |  |  |
| **Arg564Ser** | 3 | 0.38±0.12 | 6.5±2.6 | 0.65±0.10 | 1.8±0.56 | 2.8±1.1 | 0.71±0.11 |
| **Arg781His** | 3 | 0.016±0.003 | 7.1±2.0 | 0.027±0.01 | 0.20±0.04 | 3.8±1.3 | 0.062±0.01 |

Two novel frameshift variants were detected in our patient cohort: p.Ile153Tyrfs*6 and p.Cys670Serfs*14, both resulting in premature stop codons (**Table 1**). The p.Ile153Tyrfs*6 mutation results in early truncation of the ELAC2 protein, eliminating all functional motifs, and was therefore considered *a priori* as loss of function; expression and assay of this mutant was not attempted. On the other hand, p.Cys670Serfs*14 occurs closer to the carboxy terminus. We therefore tested p.Cys670Ser*14 for enzymatic activity. This mutant expressed poorly, however, and displayed no detectable enzyme activity, confirming the functional importance of domains beyond the truncation, including motif V and other functional elements (**Figure S3**). Taken together, the analysis of recombinant mutant proteins indicates that the detected *ELAC2* variants impair the RNase Z activity, consistent with pathogenicity. However, due certain limitations of the *in vitro* assay used (e.g. precursor substrate specificity or the use of substrates without post-transcriptional nucleotide modifications), we performed other functional studies fully support the pathogenic nature of the detected variants.

### Analysis of mtRNA processing in primary fibroblasts from affected individuals

In the polycistronic transcripts produced through transcription of mtDNA, the two mt-rRNAs and most mt-mRNAs are punctuated by one or more mt-tRNAs. As shown previously, impairment in the ELAC2 endonucleolytic activity results in the presence of 3’ unprocessed mt-tRNAs, containing mt-rRNA or mt-mRNA extensions [15, 36]. To analyze the levels of unprocessed mt-tRNAs resulting from impaired RNase Z activity of ELAC2, we used northern blotting with RNA samples isolated from fibroblasts of all patients that harbored novel *ELAC2* mutations. P5, P11 and P13 were excluded from this analysis as these three individuals harbored the previously characterized c.460T>C (p.Phe154Leu) variant in homozygosity [15]. In RNA samples from P1-P4, P6-P10 and P12, we found substantially increased amounts of 3′ end unprocessed mt-tRNA precursors at the cleavage sites of mt-tRNA^Val^-16S rRNA, mt-tRNA^Met^-ND2 (**Figure 3**) and mt-tRNA^Leu(UUR)^-ND1 (**Figure S2**) as compared to the control cell lines. This analysis further showed that RNA samples from the patients that were compound heterozygous for c.1690C>A (p.Arg564Ser) or c.2039C>T (p.Ala680Val) (P6 or P7, respectively) – the two mutations that exhibited little impairment with the pre-mt-tRNA^Leu(UUR)^ substrate in the *in vitro* experiments (**Figure 2b**) - accumulated unprocessed mt-tRNA^Leu(UUR)^-ND1 junctions. This latter result confirms the causal role of the p.Arg564Ser and p.Ala680Val substitutions.

**Figure 3.**
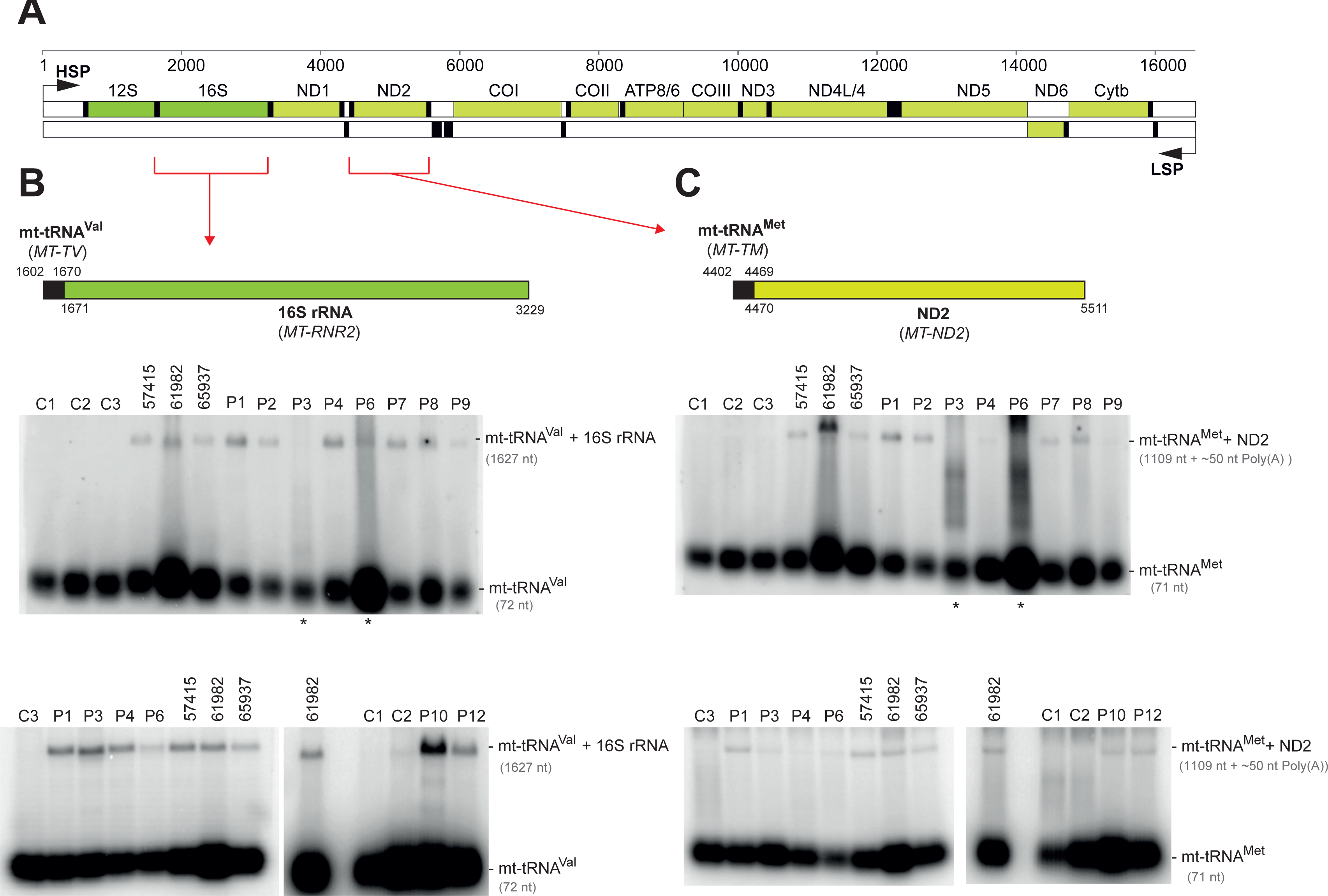
Analysis of unprocessed mitochondrial tRNA-mRNA intermediates. **(A)** Linear genetic map of mtDNA (numbering according to RefSeq accession number J01415) indicating mt-rRNA (green), mt-mRNA (olive) and mt-tRNA (black). Non-coding sequences in white. The mt-tRNA^Val^-16S rRNA and mt-tRNA^Met^-ND2 mRNA junctions are indicated by red brackets. LSP – Light strand promoter. HSP – Heavy strand promoter. **(B)** Northern blot processing analysis of the mt-tRNA^Val^-16S rRNA junction in total RNA samples of control fibroblasts (C1-C3), fibroblasts from the previously published cases (57415, 61982, 65937)[15] and fibroblasts from the patients harboring novel ELAC2 mutations (P1-4, P6-9 and P12). **(C)** Northern blot analysis of the processing of the mt-tRNA^Met^-ND2 mRNA junction. Samples as per (B). Asterisks indicate partially degraded RNA samples that were reanalysed in a different blot and presented in the same panel.

Assessing the clinical significance of sequence variants that may alter splicing, especially in the context of tissue-specific disease manifestation, can be challenging [37]. In four patients from our cohort, P2, P3, P6 and P8, compound heterozygous splice-site mutations were present. The analysis of the processing of mtRNA of these four patients revealed accumulation of unprocessed mitochondrial tRNA-rRNA or tRNA-mRNA junctions, providing evidence for the disruptive nature of the detected splice variants. Taken together, the observed defect in the processing of mt-tRNA junctions in the patient-derived cell lines provides additional evidence for the pathogenicity of all detected, novel missense, frameshift and splice site *ELAC2* variants.

### In silico characterization of mutations in ELAC2

Having established the damaging nature of the detected *ELAC2* missense mutations both *in vitro* and in living cells, we set out to develop a rationale for the observed biochemical impairment, as summarized in **Table S1**. To this end, we modelled all 13 substitutions (3 published previously and 10 novel) into the structure of *S. cerevisiae* RNase Z (Trz1) (PDB 5MTZ [30]). Analysis of the ELAC2 structure indicates that the amino and carboxy domains arose from a duplication, with the active site being retained in the carboxy domain and the flexible arm being preserved in the amino domain [38]. The amino and carboxy domains are tethered by a flexible linker (**Figure S3**). Substitutions at Arg68, Phe154, Gln388 are found in the amino domain, Leu422 and Leu423 are in the linker domain (flexible tether), whereas Pro493, Thr520, Arg564, Lys660, Arg680, Tyr729, His749, Arg781 are located in the carboxy domain of ELAC2 (**Figure 4** and **Figure S3**).

**Figure 4.**
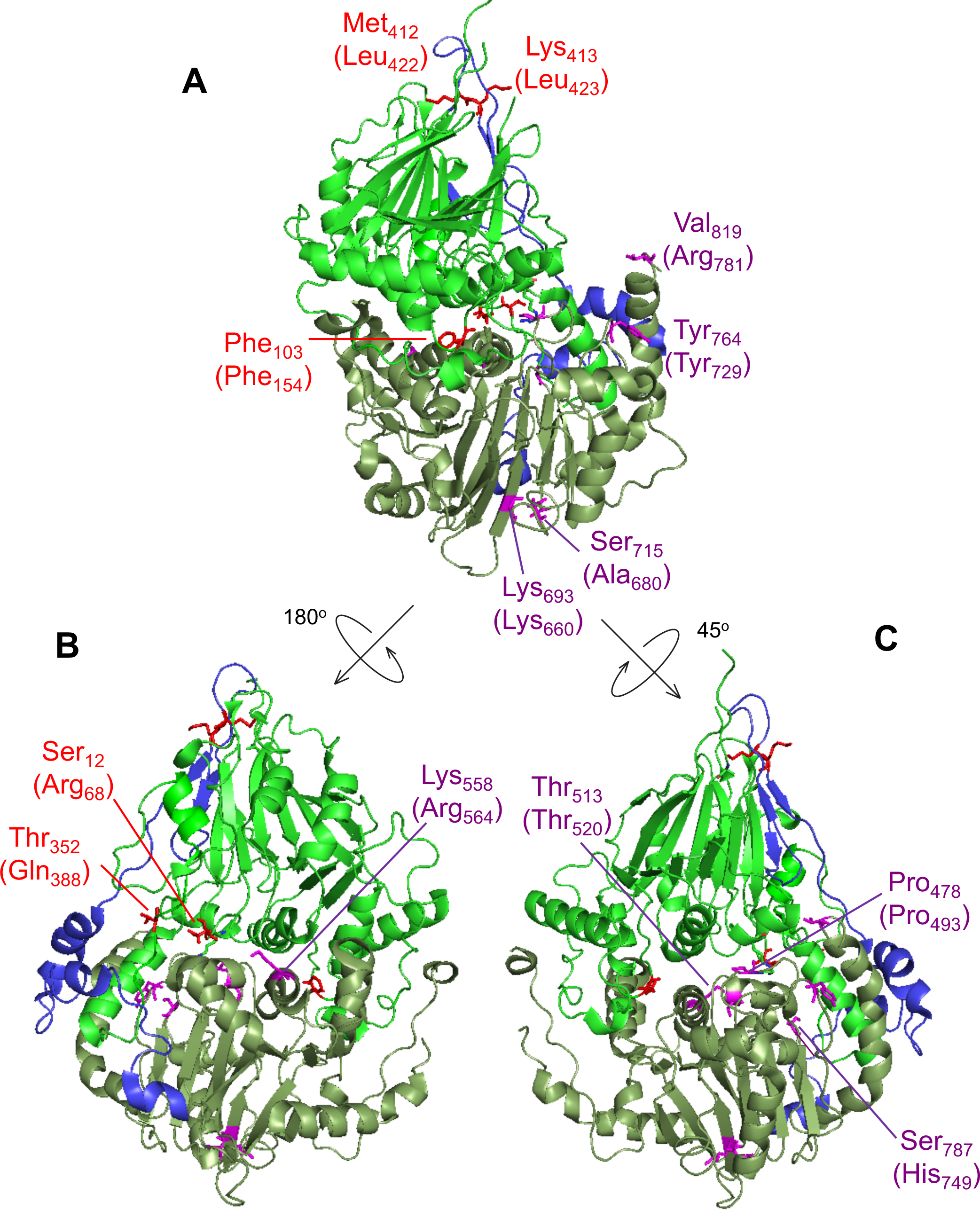
ELAC2 substitutions mapped on the structure of *Saccharomyces cerevisiae RNase Z*. The structure of *S. cerevisiae* RNase Z (Trz1, PDB#5MTZ) [30] is shown in cartoon using PyMol. The amino domain, inter-domain linker and carboxy domain are colored green, blue and pale green, respectively. Three views are shown to effectively visualize all the substitutions. All 13 ELAC2 substitutions (3 published previously [15], and 10 novel) are shown in all three views. Residues are labelled with *S. cerevisiae* RNase Z numbers and the numbers in brackets are for the *H. sapiens* ELAC2 residues. Some residues are not conserved between *S. cerevisiae* and *H. sapiens* RNase Z. Residues Arg68, Phe154, Gln388 localised in the amino domain are marked in red; Leu422 and Leu423 are in linker (also marked in red); Pro493, Thr520, Arg564, Lys660, Arg680, Tyr729, His749, Arg781 are in the carboxy domain and indicated in purple. **(A)** View with amino domain up, carboxy domain down and linker behind with the *H. sapiens* residues Phe154, Leu422, Leu423, Lys660, Ala680, Tyr729 and Arg781 labelled. **(B)** The ELAC2 model is rotated with linker on left with the residues Arg68, Gln388 and Arg564 labelled; **(C)** ELAC2 rotated with linker on right and the residues Pro493, Thr520 and His749 labelled. Note: the residues at Arg68, Phe154 and Gln388 in *H. sapiens* ELAC2 map to the domain interface.

Using the Trz1 structure, we have modelled the residues Arg68, Phe154 and Gln388 in the amino domain in which substitutions were found associated with HCM. Modelling using the Trz1 structure is less obviously effective in cases where the residues in the *S. cerevisiae* Trz1 sequence are not conserved (**Figure S3**); in these cases, however, the overall sequence in the region is conserved, and thus structural relationships of inferred secondary structure elements may be conserved. All three residues are located close to the domain interface. The modelling of Arg68 and Gln388 suggests the substitutions at these sites could disrupt the structure and folding of ELAC2 through their indirect effects on a number of polar contacts across the amino- and carboxy domain interface (**Figure S4**). Phe154 is conserved and aligns with Trz1 Phe103 (**Figure S3**). Across the N- and C-domain interface Phe154 closely approaches the motif II region, suggesting that reducing the size of the hydrophobic side chain in the case of p.Phe154Leu could affect packing of a region which is critical for metal ion binding and catalysis (**Figure S5**).

Contiguous residues Leu422 and Leu423 are located in the linker domain of the human ELAC2 at the apex of the loop between two twisted beta sheets in the amino domain. Substitutions at these positions may affect the structure due to subtle changes in regional hydrophobicity (**Figure S6**).

We next set out to model residues in the carboxy domain (Pro493, Thr520, Arg564, Lys660, Ala680, Tyr729, His749 and Arg781). All of these but two could be effectively modelled using Trz1. However, it was advantageous to model Pro493 and Tyr729 using the *Bacillus subtilis* (Bsu) enzyme-substrate complex structure (PDB 4GCW, [31]). The conserved residue Pro493 is close to Lys495 and similarly, Tyr729 is next to Arg728; equivalent residues in *Bacillus subtilis* RNase Z Lys15 and Arg273, respectively, make polar contacts with the substrate on both sides of the acceptor stem (**Figure 5**), as effectively modelled using the *Bacillus subtilis* structure. The model suggests that Pro493Leu impairs RNase Z activity by changing the fold of the PxKxRN loop, interfering with the polar contact between Lys495 and the backbone of substrate nucleotides +1 and +2 on the 5’ side of the tRNA acceptor stem. Correspondingly, substitution of Tyr729 with Cys could alter the fold of the motif V loop, interfering with contacts between the positively charged side chain of Arg728 and the substrate in the backbone of nucleotides 71, 72 and 73 on the 3’ side of the tRNA acceptor stem of the pre-tRNA substrate. In the wild type enzyme, the two basic residues, Lys495 and Arg728, clamp the substrate close to the scissile bond from both sides (**Figure 5**). This interpretation of *in silico* modelling is consistent with the kinetic experiments, in which the p.Pro493Leu and p.Tyr729Cys mutants display the greatest impairment factors of any patient-related substitutions analysed so far.

**Figure 5.**
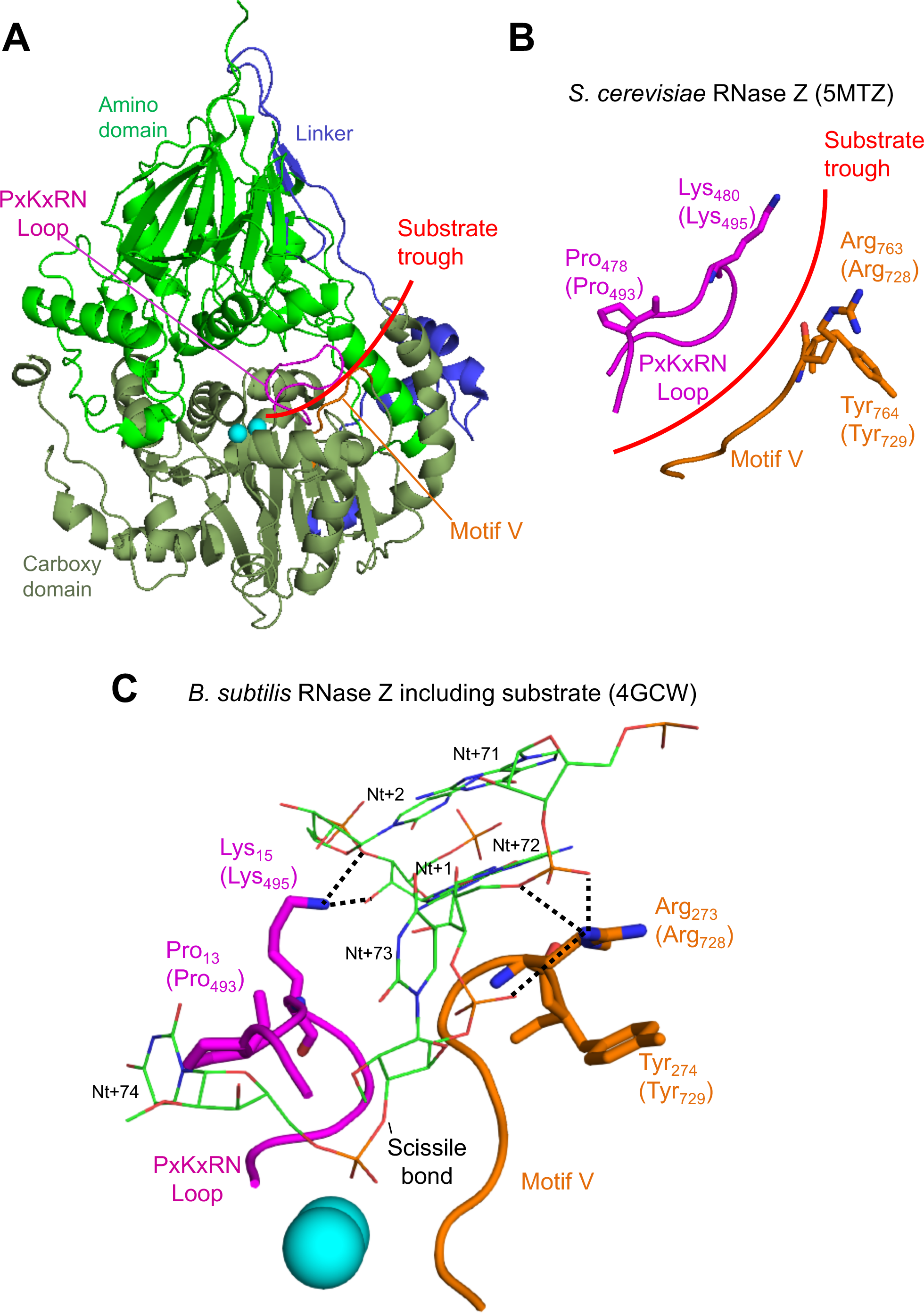
*In silico* analysis of disease-related ELAC2 substitutions p.Pro493Leu and p.Tyr729Cys. **(A)** Structure overview of *S. cerevisiae* Trz1. Amino and carboxy domains and linker are colored as in previous figures. Curved arrow indicates the presumed substrate trough. Metal ions which mark the active site are shown as blue spheres. PxKxRN and Motif V loops are shown in magenta and orange, respectively. **(B)** Detailed view of PxKxRN loop (magenta) and Motif V loop (orange) in *S. cerevisiae* Trz1. The conserved basic residues Lys in the PxKxRN loop and Arg in the Motif V loop are shown as sticks, with *H. sapiens* equivalent residues given in brackets. Red arc illustrates path of substrate acceptor stem and 3’ trailer through the presumed substrate trough. **(C)** Detailed view of PxKxRN loop (magenta) and Motif V loop (orange) in *B. subtillis* with pre-tRNA substrate (Note: The *B. subtillis* structure displays similar folds and relative orientations, validating its use for modelling ELAC2 (RNase Z^L^) structure including the carboxy domain and the active site. The *B. subtilis* RNase Z structure 4GCW [31] is the only available structure of an enzyme – pre-tRNA substrate complex). The pre-tRNA substrate acceptor stem is clamped by polar contacts with K in the PxKxRN loop (Lys495 in *H. sapiens*, Lys480 in *S. cerevisiae*, Lys15 in *B. subtilis*) and Arg in the Motif V loop (Arg728 in *H. sapiens*, Arg763 in *S. cerevisiae*, Arg273 in *B. subtilis*), illustrated with bold dashed lines. A polar substrate acceptor stem clamp consists of Lys15 contacts with 2’ and 3’ O’s of ribose +1 and Arg273 contacts with two backbone phosphate O’s on nt 72 and one on nt 73. Counterintuitively, the basic R-groups that form the substrate clamp do not point toward each other; both are oriented toward the right, but two polar contacts extend to the right from Lys15 while three polar contacts by Arg273 extend to the left toward substrate.

Thr520 (Thr513 in the Trz1 structure) is a highly conserved motif I residue. A charge relay system links the motif I aspartate (Asp515) with the first histidine of motif II (His546). The model implies that subtle regional changes impair catalysis, transmitted to motif I (**Figure S7**).

Arg564 is not conserved, and is located in the long, overall poorly conserved region between motifs II and III, with no significant polar or hydrophobic contacts in this region of the Trz1 structure (**Figure S3**). The mutated residue is reasonably close to motif II, implying that subtle regional changes may impair catalysis when transmitted to this motif.

The side chain of conserved Lys660 (Lys693 in the Trz1 structure) found in **β**24 is predicted to make polar contacts with the backbone of the preceding residues within 12 residues of the motif IV aspartate (**Figure S4** and **S8**). Loss of these side chain-specific polar contacts with the p.Lys660Ala substitution could therefore affect the position of Asp666, the motif IV aspartate, which is critical for metal ion binding and catalysis (**Figure S8B**). Similarly, Ala680 (Ser715 in the Trz1 structure) located below the base of **β**25 could, when replaced by the bulkier valine in the case of the Ala680Val mutation, indirectly affect the location of the Motif IV aspartate (**Figure S8B**). Finally, His749 (Ser787 in the Trz1 structure), is predicted to be contiguous with the aspartate in the AxD loop and substitution at this position could perturb that residue. The AxD loop is conserved and the aspartate in this loop makes polar contacts with the backbone of a conserved leucine at the start and a conserved asparagine at the end of the PxKxRN loop (**Figure S8B**; cf [39]).

The Arg781 residue is present on a long C-terminal α-helix. Although this residue is not conserved, the region where it is found is generally highly polar. In metazoan ELAC2 enzymes this region consists of frequently interspersed acidic and basic residues while in *Saccharomyces cerevisia*e it is principally acidic (**Figure S3**). In *S. cerevisiae* RNase Z, the long α-helix is curved and approaches the predicted location where the substrate acceptor stem is clamped by polar contacts between Lys495 of the PxKxRN loop and nt +1-2 of the pre-tRNA substrate and between Arg728 of the motif V loop and nt 71-72-73 of the substrate (**Figure 5** and **Figure S9**). The region where Arg781 is found could thus modulate both substrate binding and catalysis, consistent with impairment of the p.Arg781His mutant which arises from the combination of a reduction in *k*_cat_ and increased *K*_M_, (**Table 3** and **Table 4**).

## Discussion

The mitochondrial genome encodes key subunits of the OXPHOS system and RNA components needed for mitochondrial translation, with nuclear genes encoding the proteins responsible for mtDNA transcription, post-transcriptional RNA processing and translation. Recent years have seen a rapid development in our understanding of these machineries both in human health and in disease state [40]. Dysfunction of mitochondrial gene expression, caused by mutations in either the mitochondrial or nuclear genomes, is associated with a diverse group of human disorders characterized by impaired mitochondrial respiration. Within this group, an increasing number of mutations have been identified in nuclear genes involved in endonucleolytic processing of precursor mtRNA [15, 41, 42] and in mtRNA epitranscriptome shaping [43–53].

### Genotype-phenotype correlation

Clinical syndromes associated with defects in mtRNA metabolism are characterized by the variable combination of encephalopathy, myopathy, sideroblastic anemia, cardiomyopathy, hearing loss, optic atrophy, and renal or liver dysfunction [54, 55]. Hypertrophic cardiomyopathy and lactic acidosis are frequent presentations in some mitochondrial diseases related to dysfunctional mt-tRNA maturation, such as those caused by biallelic variants in *MTO1*, *GTPBP3*, *AARS2* and *RARS2* [45, 47, 56, 57]. With multiple novel variants in *ELAC2,* our study further underscores HCM as a manifestation of dysfunctional mitochondrial gene expression. Moreover, it indicates a strong relationship between a confirmed molecular diagnosis of *ELAC2*-related mitochondrial disease and the key clinical phenotypes (hypertrophic cardiomyopathy, multiple respiratory chain defects and lactic acidosis) for the majority of variants detected. However, one case (P2), who harbors a c.1423+1G>A *ELAC2* variant affecting a consensus splice site (in compound heterozygosity with a truncating c.2009del; p.Cys670Serfs*14 variant), was the only patient in our cohort who did not present with cardiomyopathy. Interestingly, five patients from an inbred, consanguineous Arabic family harboring another homozygous *ELAC2* mutation involving the same consensus splice site (homozygous c.1423+2T>A) predominantly presented with intellectual disability without prominent cardiac involvement. Both of these variants affect the same consensus donor sequence (exon 15), and it would be interesting to further explore the features of pre-mRNA splice site selection of this particular donor site in cardiac tissue. Interestingly, mtDNA depletion was detected in the hypertrophic heart and not the skeletal muscle of P7 [32]. To the best of our knowledge, quantitative abnormalities of the mtDNA has so far never been associated with defects in mtRNA processing, and additional observations are necessary to confirm this association.

### Enzymatic mechanism of ELAC2

*In vitro* analysis of the *ELAC2* mutants allowed for a better understanding of the enzymology of this mitochondrial RNase Z. In particular, the p.Pro493Leu and p.Tyr729Cys substitutions most severely impaired enzyme function, resulting in ~1% of WT enzyme activity. No mutations were found which interfere directly with metal ion binding or catalysis, although numerous such substitutions, constructed by site directed mutagenesis, greatly impair catalysis, between 500 – 10,000-fold relative to WT [58, 59]. Relatively severe impairment by substitutions of the Pro493 and Tyr729 residues probably arises from their proximity in loops to the charged residues that clamp the substrate acceptor stem close to the scissile bond. Replacement of proline with leucine, although similar in hydrophobicity, could alter the path of the PxKxRN loop, affecting the position and orientation of Lys495. With substitution of Cys for Tyr729 in the motif V loop, the reduced side chain size could affect the path of the loop, including position and orientation of neighboring Arg728, the arginine residue which makes critical polar contacts with the acceptor stem of substrate. In this way the most severe impairment arises from proximity to conserved residues with established functions. In this context, our study of naturally occurring pathogenic mutations provides further insights into the enzymatic mechanism of ELAC2.

### Missense variant associated with prostate cancer impairs ELAC2 enzymatic activity on mitochondrial substrates

The p.Ala541Thr and p.Arg781His ELAC2 substitutions were first characterized as variants in a pedigree in Utah displaying early-onset prostate and other cancers [60]. The appearance of the frequent *ELAC2* polymorphism p.Ala541Thr in P1 of this study appears to be coincidental and without biological significance, as p.Pro493Leu and p.Arg68Trp have been documented here as responsible for pathogenicity. The independent reappearance of the missense substitution p.Arg781His in P3, P4 and P10, which is exceedingly rare in the general population (MAF= 0.0005165 in gnomAD), is here shown to be significant in the context of mitochondrial tRNA metabolism. In previously published data, no differences in catalysis were observed between wild type and prostate cancer-associated mutants of ELAC2 (the missense substitution and two much more frequent polymorphisms) using nuclear-encoded pre-tRNA substrates [61, 62] (Yan and Levinger, unpublished observations). The p.Arg781His substitution, which here re-emerged in three described cases of HCM (Patients 3, 4 and 10), clearly impairs processing of mitochondrial pre-tRNA substrates both *in vitro* and in living cells, suggesting that the associated phenotypes (possibly including prostate cancer susceptibility) are mitochondrially-based.

Catalytic efficiencies with WT enzyme are generally lower using mitochondrial substrates than with nuclear-encoded substrates and the lower catalytic efficiency is more pronounced with mt-tRNA^Ile^ than with mt-tRNA^Leu(UUR)^. Two levels of catalytic activity were thus observed, between nuclear vs. mitochondrial and further among different mitochondrial tRNAs [28, 29]. To explain the first level distinction, mitochondrial tRNAs generally have a weaker and non-canonical secondary and tertiary structure (reviewed in [63]). For example, mt-tRNA^Leu(UUR)^ displays a heterogeneous secondary structure and among mitochondrial tRNAs, it is closer to canonical than mt-tRNA^Ile^, which displays AU rich stems including an A/C mismatch in the T-stem [64]. Wild type substrate structures that reduce wild type ELAC2 catalytic efficiencies may thus lead to more pronounced impairment with missense substitutions which otherwise were not observed with more canonical nuclear encoded substrates.

## Conclusions

Pathogenic variants in *ELAC2* impair the RNase Z activity of this critical mitochondrial enzyme. Decreased ELAC2 activity leads to a disturbance of the proper mitochondrial gene expression by increasing the amounts of incorrectly processed mtRNA. The consequence of perturbed ELAC2 function is manifested by multiple mitochondrial respiratory chain deficiencies, hypertrophic cardiomyopathy and lactic acidosis. Therefore, the *ELAC2* gene should be included in gene panel to screen infantile-onset cases of hypertrophic cardiomyopathy patients. Association between the c.2342G>A (p.Arg781His) *ELAC2* variant and prostate cancer as a consequence of impaired mitochondrial RNase Z activity could indicate a functional link between tumorigenesis and mitochondrial RNA metabolism.

## Acknowledgments

C.A.P., P.R-G. and M.M. were supported by Medical Research Council, UK (MC_U105697135 and MC_UU_00015/4). P. R.-G. is supported by Fundação para a Ciência e a Tecnologia (PD/BD/105750/2014). M.S., M.P., K-G. P. and L.L. were supported by the grant R15GM101620 from the NIH. L.L. is a member of the graduate programs in Biochemistry and Molecular and Cellular Biology, The City University of New York. R.M. and R.W.T. are supported by the Wellcome Centre for Mitochondrial Research (203105/Z/16/Z), the MRC Centre for Neuromuscular Diseases (G0601943), Newcastle University Centre for Ageing and Vitality (supported by the Biotechnology and Biological Sciences Research Council and Medical Research Council (G016354/1)), the UK NIHR Biomedical Research Centre in Age and Age Related Diseases award to the Newcastle upon Tyne Hospitals NHS Foundation, the MRC/ESPRC Newcastle Molecular Pathology Node, the Lily Foundation and the UK National Health Service Highly Specialised Service for Rare Mitochondrial Disorders. C.L.A. was supported by a National Institute for Health Research (NIHR) doctoral fellowship (NIHR-HCS-D12-03-04). R.H. is a Wellcome Investigator (109915/Z/15/Z), who receives support from the Medical Research Council (UK) (MR/N025431/1), the European Research Council (309548), the Wellcome Trust Pathfinder Scheme (201064/Z/16/Z) and the Newton Fund (UK/Turkey, MR/N027302/1). A.A. receives funding for a PhD studentship from the Kuwait Civil Service Commission under the approval of the Kuwait Ministry of Health. D.G. was supported by the Telethon Foundation (Grant GGP15041); the Pierfranco and Luisa Mariani Foundation; the E-Rare project GENOMIT. H.P. was supported by the German Bundesministerium für Bildung und Forschung (BMBF) and Horizon2020 through the E-Rare project GENOMIT (01GM1603 and 01GM1207), through the German Network for mitochondrial disorders (mitoNET, 01GM1113C) and the EU Horizon2020 Collaborative Research Project SOUND (633974). The “Cell lines and DNA Bank of Genetic Movement Disorders and Mitochondrial Diseases” of the Telethon Network of Genetic Biobanks (grant GTB12001J) and the EuroBioBank Network supplied biological specimens. E.B., R.C. M.D.N and D.V. are supported by the Italian Ministry of Health Ricerca Corrente. S.R. receives grant funding from Great Ormond Street Hospital Children’s Charity and the Lily Foundation.

## Authors Contributions

**Conceptualization**: M.M., L.L, H.P., R.W.T;

**Methodology**: S.M., L.L., M.M., C.A.P.;

**Validation**: M.M., L.L, H.P. R.W.T.;

**Formal analysis**: M.S., C.A.P., R.K., B.L., M.M., L.L, H.P. R.W.T, D.G.;

**Investigation**: M.S., C.A.P., P.R-G., R.K., D.G., F.I., E.L., M.P., K-G. P., A.A. H.H.A-B,

B.A. C.A.L, D.B., M.D.N, D.V., S.R.;

**Resources**: M.A.D, S.S., P.B., M.A., R.C., A.G., E.H., D.G., S.R., R.H.;

**Writing—original draft:** M.M., L.L, R.W.T., R.M.

**Writing—review & editing**: All authors;

**Visualization**: P.R-G., M.S., C.A.P, L.L., M.M.;

**Supervision**: M.M., L.L, H.P. R.W.T, D.G., E.B., E.H.

**Project administration**: M.M., L.L, H.P. R.W.T, D.G., E.B., E.H.

**Funding acquisition:** M.M., L.L, H.P. R.W.T, D.G., E.B., R.H.

